# The CRP-family transcriptional regulator DevH regulates expression of heterocyst-specific genes at the later stage of differentiation in the cyanobacterium *Anabaena* sp. strain PCC 7120

**DOI:** 10.1101/2020.04.22.054668

**Authors:** Yohei Kurio, Yosuke Koike, Yu Kanesaki, Satoru Watanabe, Shigeki Ehira

**Affiliations:** Department of Biological Sciences, Graduate School of Science, Tokyo Metropolitan University, Tokyo, Japan; NODAI Genome Research Center, Tokyo University of Agriculture, Tokyo, Japan; Department of Bioscience, Tokyo University of Agriculture, Tokyo, Japan

**Keywords:** *Anabaena*, heterocyst differentiation, microoxic intracellular environment, nitrogenase, transcriptional regulation

## Abstract

Heterocysts are terminally differentiated cells of filamentous cyanobacteria, which are specialized for nitrogen fixation. Because nitrogenase, an enzyme for nitrogen fixation, is easily inactivated by oxygen, the intracellular environment of heterocysts is kept microoxic. In heterocysts, the oxygen-evolving photosystem II is inactivated, a heterocyst-specific envelope with an outer polysaccharide layer and an inner glycolipid layer is formed to limit oxygen entry, and oxygen consumption is activated. Heterocyst differentiation, which is accompanied by drastic morphological and physiological changes, requires strictly controlled gene expression systems. Here, we investigated the functions of a CRP-family transcriptional regulator, DevH, in the process of heterocyst differentiation. A *devH*-knockdown strain, devHKD, was created by replacing the original promoter with the *gifA* promoter, which is repressed during heterocyst differentiation. Although devHKD formed morphologically distinct cells with the heterocyst envelope polysaccharide layer, it was unable to grow diazotrophically. Genes involved in construction of the microoxic environment, such as *cox* operons and the *hgl* island, were not upregulated in devHKD. Moreover, expression of the *nif* gene cluster was completely abolished. Even under anaerobic conditions, the *nif* gene cluster was not induced in devHKD. Thus, DevH is necessary for the establishment of a microoxic environment and induction of nitrogenase in heterocysts.

## Introduction

Nitrogen fixation is a crucial process for sustaining living organisms on the earth by providing a nitrogen source in a bioavailable form. Cyanobacteria play an important role in various ecosystems as primary producers by conducting oxygen-evolving photosynthesis and nitrogen fixation. Some filamentous cyanobacteria such as *Anabaena* sp. strain PCC 7120 (hereafter *Anabaena* PCC 7120) form specialized cells called heterocysts for nitrogen fixation. Filaments of *Anabaena* PCC 7120 are composed of a few hundred vegetative cells in the presence of combined nitrogen in the medium, and 5% to 10% of vegetative cells differentiate into heterocysts upon depletion of combined nitrogen (Flores *et al*., 2019). Heterocysts maintain the microoxic intracellular environment to protect O_2_-labile nitrogenase through inactivating O_2_-producing photosystem II and activating O_2_ consumption by terminal oxidases and flavodiiron proteins (Valladares *et al*., 2007; Ermakova *et al*., 2014). In addition, a heterocyst-specific envelope, which consists of layers of polysaccharides and glycolipids, is formed outside of the cell wall to limit O_2_ entry into heterocysts (Murry and Wolk, 1989). Thus, heterocyst differentiation is accompanied by changes in cellular physiology and morphology, which are caused by the spatiotemporal regulation of gene expression (Ehira *et al*., 2003; Ehira and Ohmori, 2006a; Flaherty *et al*., 2011; Mitschke *et al*., 2011).

The intracellular level of 2-oxoglutarate (2-OG) represents cellular nitrogen status (Muro-Pastor *et al*., 2001) and its increase triggers heterocyst differentiation (Laurent *et al*., 2005). 2-OG directly binds to and activates a cAMP receptor protein (CRP)-family transcriptional regulator, NtcA, which is a global regulator of nitrogen and carbon metabolism in cyanobacteria (Vázquez-Bermúdez *et al*., 2002; Tanigawa *et al*., 2002). NtcA induces the expression of a response regulator NrrA, which is necessary for the full induction of *hetR* (Muro-Pastor *et al*., 2006; Ehira and Ohmori, 2006b). HetR is a master regulator of heterocyst differentiation and is essential and sufficient for this process (Buikema and Haselkorn, 1991; Buikema and Haselkorn, 2001). HetR directly regulates the expression of *hetP* and *hetZ*, both of which are necessary for heterocyst differentiation (Zhang *et al*., 2007; Higa and Callahan, 2010; Videau *et al*., 2018), and cells highly expressing *hetR* turn into heterocysts (Asai *et al*., 2009; Fukushima and Ehira, 2018). In the cells committed to differentiation, the expression of clusters of genes that are involved in synthesis of the heterocyst-specific envelope’s polysaccharides and glycolipids, which are called the *hep* and *hgl* islands, respectively (Ehira *et al*., 2003; Huang *et al*., 2005; Fan *et al*., 2005), is induced. In addition, respiration enzymes are upregulated in heterocysts (Valladares *et al*., 2007; Ermakova *et al*., 2014), followed by the induction of nitrogenase from the *nif* gene cluster under microoxic conditions. Two sigma factors, SigC and SigE, whose expression is regulated by *hetR*, are part of the system regulating genes exclusively expressed in heterocysts, but their inactivation does not prevent heterocyst formation (Mella-Herrera *et al*., 2011; Ehira and Miyazaki, 2015). Thus, the regulatory system at the early stage of heterocyst differentiation has been analyzed in detail, but the transcriptional regulation related to changes in heterocyst morphology and physiology remains to be revealed.

Recently, a transcriptional regulator, CnfR, has been shown to regulate expression of the *nif* gene cluster in the non-heterocyst-forming filamentous cyanobacterium *Leptolyngbya boryana* (Tsujimoto *et al*., 2014). CnfR is necessary for inducing the expression of all of the *nif* genes under nitrogen-depleted and low-oxygen conditions. In the heterocyst-forming cyanobacterium *Anabaena variabilis* ATCC 29413, there are two homologs of CnfR, CnfR1 and CnfR2 (Pratte and Thiel, 2016). CnfR2, like as CnfR in *L. boryana*, regulates the expression of the *nif2* gene cluster in vegetative cells under nitrogen-depleted and low-oxygen conditions, while CnfR1 is involved in regulating the *nif1* gene cluster, which is exclusively expressed in heterocysts (Thiel and Pratte, 2014). *Anabaena* PCC 7120 possesses only one *cnfR* gene, which is a homolog of *cnfR1*. In *Anabaena* PCC 7120, *cnfR*, previously known as *patB*, is mainly expressed in heterocysts and important for nitrogen fixation, but it is unknown whether *cnfR* regulates expression of the *nif* gene cluster (Liang *et al*., 1993; Jones *et al*., 2003).

The *devH* gene has been identified as a gene upregulated during heterocyst differentiation in *Anabaena* PCC 7120 and its upregulation depends on *ntcA* and *hetR* (Hebbar and Curtis, 2000). *devH* encodes a CRP-family transcriptional regulator, and its insertional mutant strain A57 is unable to conduct nitrogen fixation aerobically. In A57, induction of the *hgl* genes, *hglB, hglC, hglD* and *hglE*, during heterocyst differentiation is lowered and the heterocyst-specific glycolipids are not synthesized, indicating that *devH* is required for formation of the heterocyst-specific glycolipid layer (Ramírez *et al*., 2005). Although A57 shows a very low level of nitrogenase activity under anaerobic conditions, the transcripts for *nifHDK*, which are the structural genes of nitrogenase, are not detected (Ramírez *et al*., 2005). These findings suggest that DevH plays an important role in regulating heterocyst differentiation, but only limited functional analysis of it has been performed. In this study, we created a *devH*-knockdown strain of *Anabaena* PCC 7120, and revealed that DevH regulates gene expression at the later stage of heterocyst differentiation. DevH was shown to be necessary for the establishment of a microoxic environment and the induction of nitrogenase in heterocysts.

## Results

### *Knockdown of the* devH *gene*

To inactivate the *devH* gene in *Anabaena* PCC 7120, we attempted to produce insertion mutants, in which antibiotic resistance cassettes were inserted at 360 bases downstream of the 5′ end of the *devH* gene. However, we were unable to obtain completely segregated clones. Hence, a knockdown mutant of *devH*, in which the *devH* promoter was replaced with the *gifA* promoter, was created (Fig. 1A). The transcription of *gifA* is repressed under nitrogen-deprived conditions by the binding of NtcA to its promoter (Galmozzi *et al*., 2010). The expression of *devH* was increased by nitrogen deprivation in the wild-type strain (WT), but in the *devH*-knockdown mutant, devHKD, the *devH* expression was the highest in the presence of ammonium and markedly decreased by nitrogen deprivation (Fig. 1B). The *devH* transcript levels at 8 and 24 h after nitrogen deprivation in devHKD were one-tenth and one-fifth of those in WT, respectively, and were lower than the lowest level in WT (0 h). devHKD grew normally in the presence of ammonium as a nitrogen source, while under diazotrophic growth conditions it did not proliferate (Fig. 1C). In devHKD, heterocysts were visually detectable, and the polysaccharide layer of the heterocyst envelope that was stained with Alcian blue was formed (Fig. 1D). Thus, devHKD is capable of initiating heterocyst differentiation, but would be incapable of completing differentiation. These phenotypes of devHKD are similar to those of the *devH* mutant strain A57 (Hebbar and Curtis, 2000; Ramírez *et al*., 2005), indicating that the DevH expression in devHKD is lowered to a level that is inadequate for its function.

**Fig. 1.**
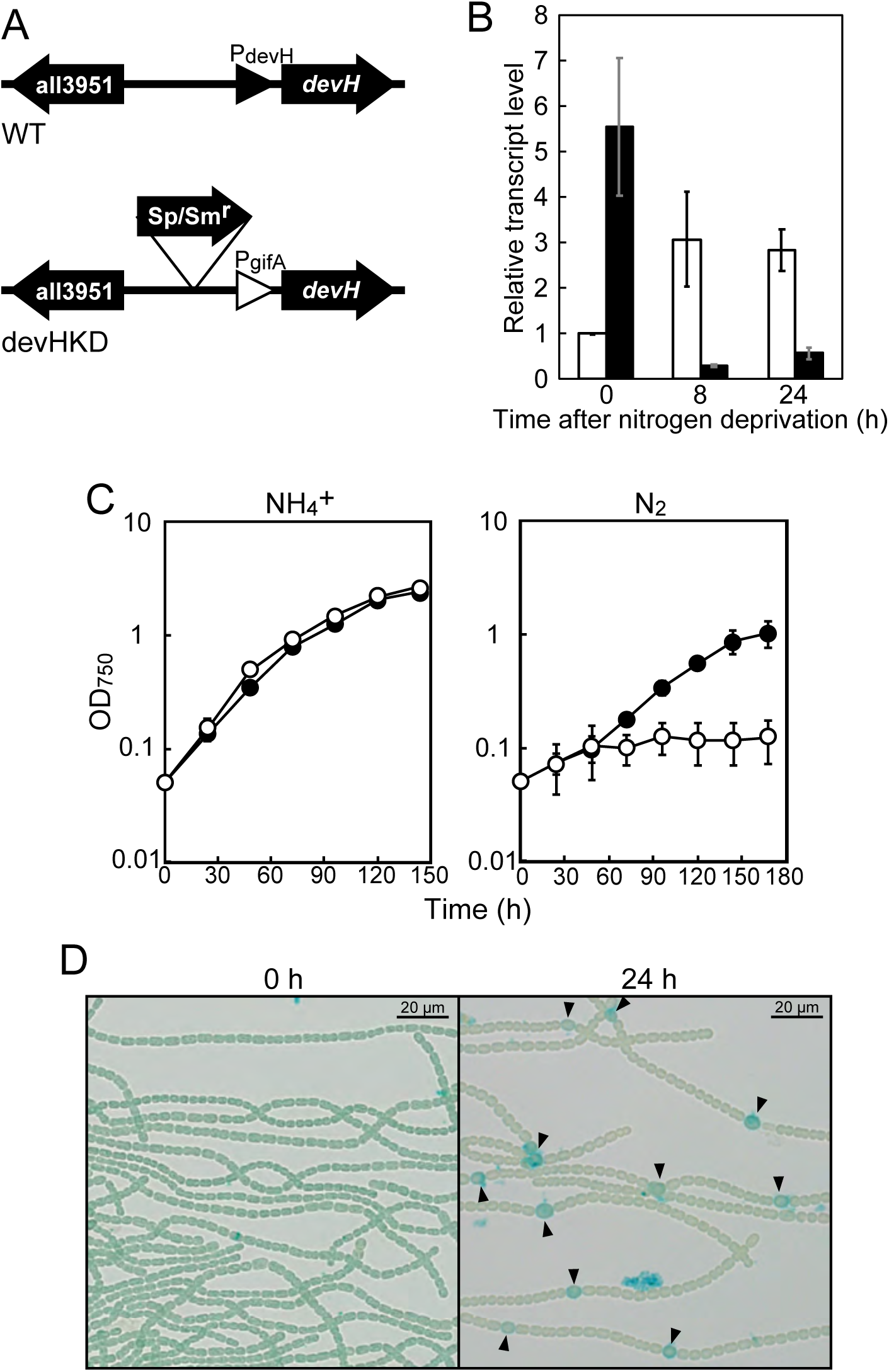
The *devH*-knockdown strain devHDK. (A) The *devH* promoter of *Anabaena* PCC 7120 (WT) was replaced with the *gifA* promoter, which is repressed during heterocyst differentiation, to construct the *devH*-knockdown strain devHKD. (B) The *devH* transcript levels during heterocyst differentiation in WT (white bars) and devHKD (black bars). (C) Growth of WT (black circles) and devHKD (white circles) on ammonium (NH_4_^+^) or dinitrogen (N_2_) as the nitrogen source. (D) Bright field images of devHKD before (0 h) and 24 h after nitrogen deprivation. Arrowheads indicate cells stained with Alcian blue. The means ± SD (error bar) of at least three independent experiments are shown.

### Identification of genes regulated by DevH

To determine the DevH regulon, RNA sequencing analysis was conducted in the WT and devHKD. DevH-regulated genes were identified by comparing the gene expression profile at 24 h after nitrogen deprivation between WT and devHKD. The gene expression profile of each strain was analyzed by RNA sequencing with two biological replicates, and genes showing decreased expression by a factor of 2 in devHKD in both experiments were identified (Table S1). The transcript levels of 135 genes were downregulated in devHKD, which included genes necessary for nitrogen fixation in heterocysts. The expression of all genes of the *nif* cluster decreased in devHKD (Fig. 2). Genes of the *hgl* island, the *cox2* and *cox3* operons, and *flv3B*, which maintain a microoxic environment in heterocysts (Valladares *et al*., 2003; Fan *et al*., 2005; Allahverdiyeva *et al*., 2013), were downregulated. In addition, the expression of *hupS* encoding uptake hydrogenase was also lowered. Genes of the *hep* island, which is required for the synthesis of the heterocyst envelope’s polysaccharides, are also indispensable for nitrogen fixation (Huang *et al*., 2005). However, expression of the *hep* island was not affected in devHKD (Fig. S1A). Changes of expression of some representative genes after nitrogen deprivation were determined by quantitative reverse-transcription PCR (qRT-PCR). The expression of all tested genes except *hepA* and *hepB* was induced between 8 and 24 h after nitrogen deprivation in WT, and the induction was abolished or suppressed in devHKD (Fig. 3). *hepA* and *hepB* were induced within 8 h in both WT and devHKD (Fig. S1B), indicating that DevH regulates gene expression at the later stage of heterocyst differentiation.

**Fig. 2.**
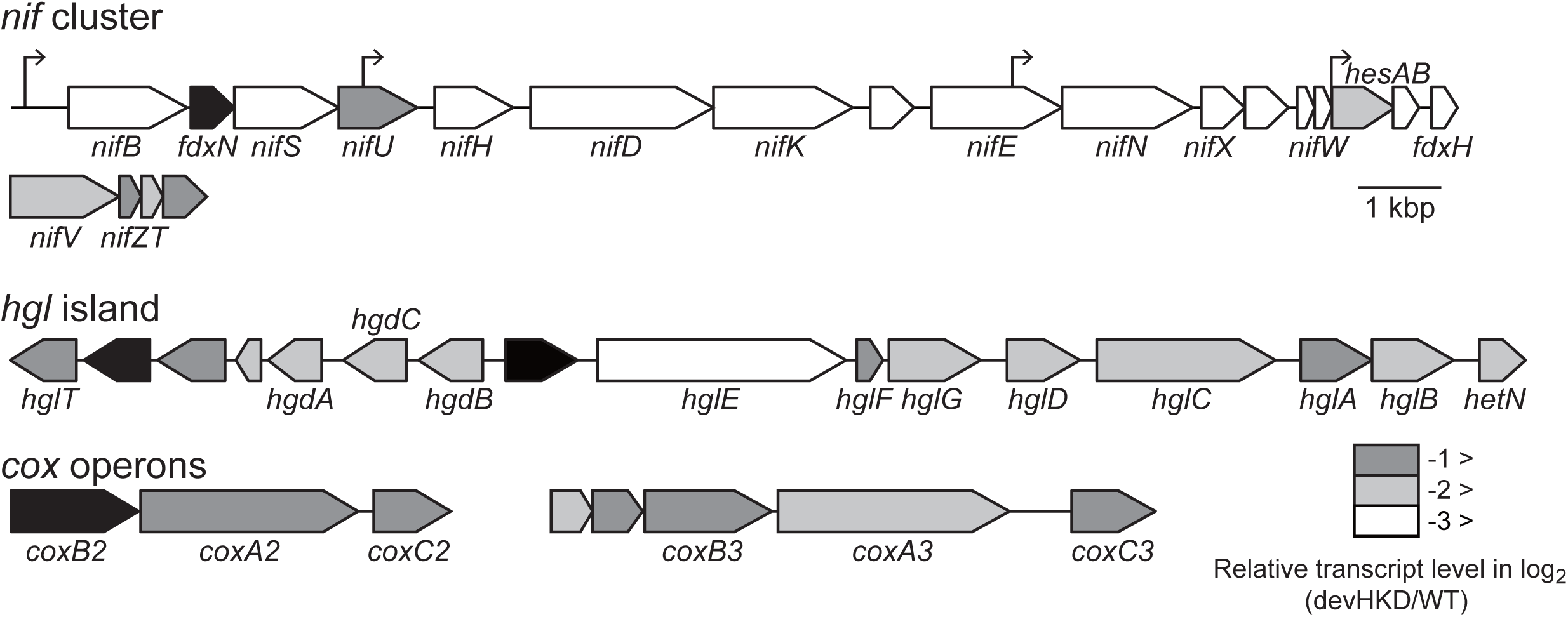
Gene clusters downregulated in devHKD. The transcript level of each gene in devHKD relative to WT, which is the mean of values of two independent RNA-sequencing analyses, is shown with a heatmap. Genes with no significant difference in expression are shown in black. The transcription start sites of the *nif* gene cluster that were identified in *A. variabilis* ATCC 29413 are shown with bent arrows.

**Fig. 3.**
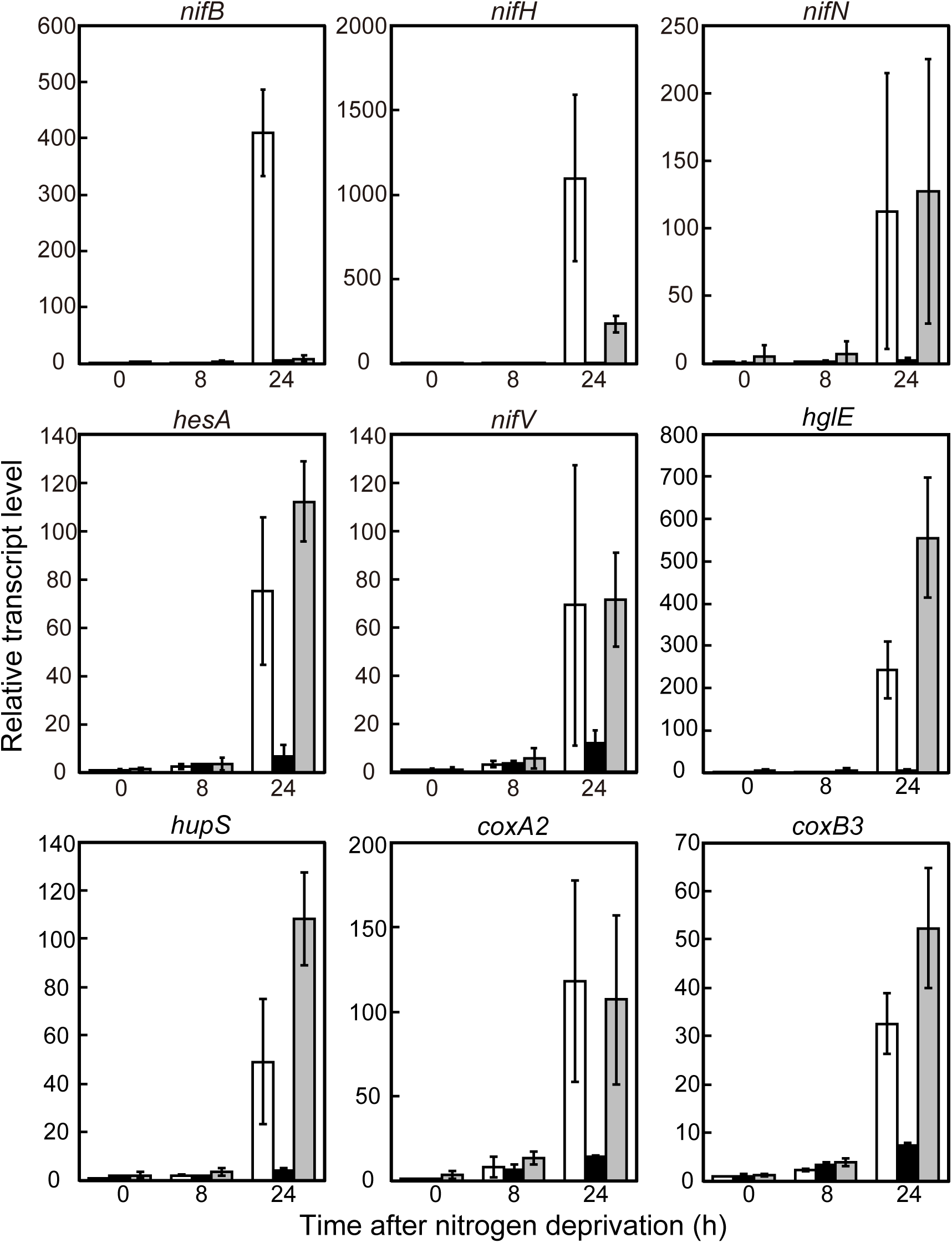
Expression of genes upregulated at the later stage of heterocyst differentiation in WT (white bars), devHKD (black bars), and a *cnfR* mutant strain (gray bars). The relative transcript levels before (0 h) and 8 or 24 h after nitrogen deprivation were determined by qRT-PCR. The qRT-PCR analysis was conducted in duplicate using three independently grown cultures. The transcript level at 0 h of WT was taken as 1.

### *Regulation of the* nif *cluster by CnfR (PatB) and DevH*

The *nif* cluster of *Anabaena* PCC 7120 is very similar to the *nif1* cluster of *A. variabilis*, both of which are exclusively expressed in heterocysts (Thiel and Pratte, 2014). There are four promoters for the *nif1* cluster: upstream of *nifB1*, within *nifU1* and *nifE1*, and upstream of *hesA1* (Fig. 2). The *nifB1* promoter is highly active, transcription from which proceeds to at least *nifE1*, and is regulated by CnfR1 (Pratte and Thiel, 2016). In *Anabaena* PCC 7120, the deletion mutant of *cnfR* (previously known as *patB*) is unable to grow under nitrogen-fixing conditions and shows a trace amount of nitrogenase activity (Jones *et al*., 2003). To clarify the roles of DevH and CnfR in the regulation of the *nif* cluster expression in *Anabaena* PCC 7120, a *cnfR* mutant, DRcnfRK, was produced. In DRcnfRK, expression of *nifB* was completely abolished, while low-level expression of *nifH* was detected (Fig. 3). Expression of *nifN, hesA*, and *nifV* as well as *hglE, hupS, coxA2*, and *coxB3* was not affected by the *cnfR* disruption. Thus, as is the case in *A. variabilis*, CnfR of *Anabaena* PCC 7120 regulates only the *nifB* promoter.

Although expression from the *nifB* promoter depends on CnfR, it has been completely abolished in devHKD. Hence, the transcript levels of *cnfR* in devHKD were determined. Expression of *cnfR* increased upon nitrogen deprivation, with the *cnfR* transcript level increasing about 25-fold after 24 h (Fig. 4A). The *cnfR* expression also increased upon nitrogen deprivation in devHKD, but the *cnfR* transcript level at 24 h in devHKD was half that in WT. We conducted gel mobility shift assays with purified DevH protein and a DNA probe of the *cnfR* promoter region. Because DevH protein was insoluble when it was expressed in *Escherichia coli*, a solubilizing tag ProS2 was fused to the N-terminus of DevH. DevH recombinant proteins bound to the *cnfR* promoter region in a sequence-specific manner (Fig. 4B), while they did not specifically interact with the *nifB* promoter (Fig. S2). DevH is involved in regulating *cnfR* expression, but the *cnfR* expression is inducible without DevH.

**Fig. 4.**
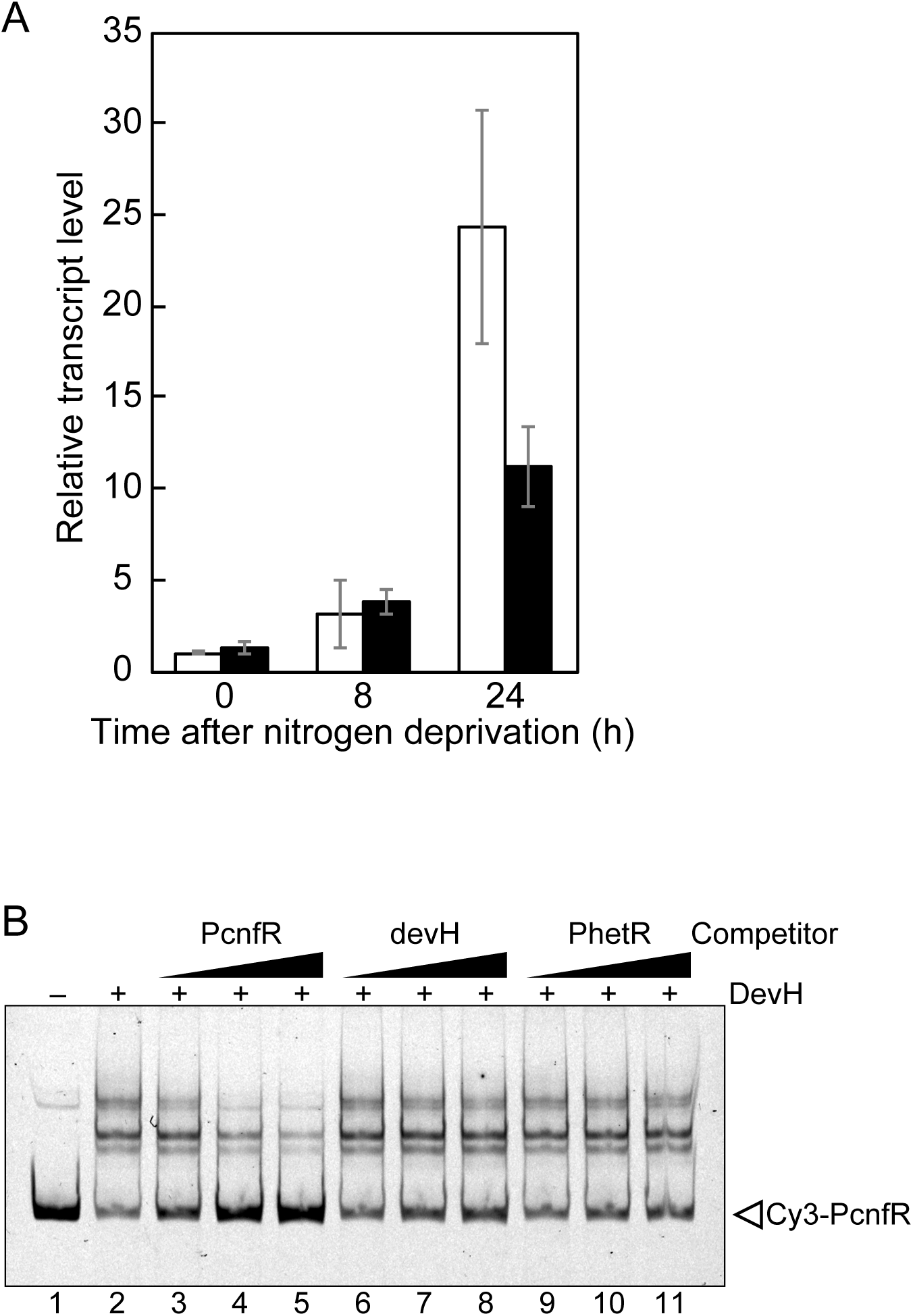
Regulation of the *cnfR* expression by DevH. (A) The *cnfR* transcript levels were determined by qRT-PCR in WT (white bars) and devHKD (black bars) as described in Fig. 3. (B) Gel mobility shift assay with recombinant DevH proteins and the *cnfR* promoter region (PcnfR). Cy3-labeled PcnfR (3 nM) was mixed with DevH (0.3 µM), and then mixtures were subjected to electrophoresis. Unlabeled probes of PcnfR (lanes 3 to 5), the devH-coding region (devH) (lanes 6 to 8), and the *hetR* promoter region (PhetR) (lanes 9 to 11) were added at concentrations of 2.5 nM (lanes 3, 6, and 9), 20 nM (lanes 4, 7, and 10), and 50 nM (lanes 5, 8, and 11). Lane 1, DevH was not added.

In addition to *cnfR*, DevH was likely to directly regulate the expression of *hglE* and itself. Four transcription start sites (TSSs) were detected for *hglE* (Imashimizu *et al*., 2005). Two probes, PhglE1 and PhglE2, which include two proximal TSSs and two distal TSSs, respectively, were prepared for gel mobility shift assays. DevH specifically bound to both probes (Fig. 5A). The interaction of DevH and the *devH* promoter region was reported by Ramírez et al. (2005), although the specific data were not published. We confirmed that DevH bound to the *devH* promoter, suggesting the presence of autoregulation of DevH (Fig. 5B).

**Fig. 5.**
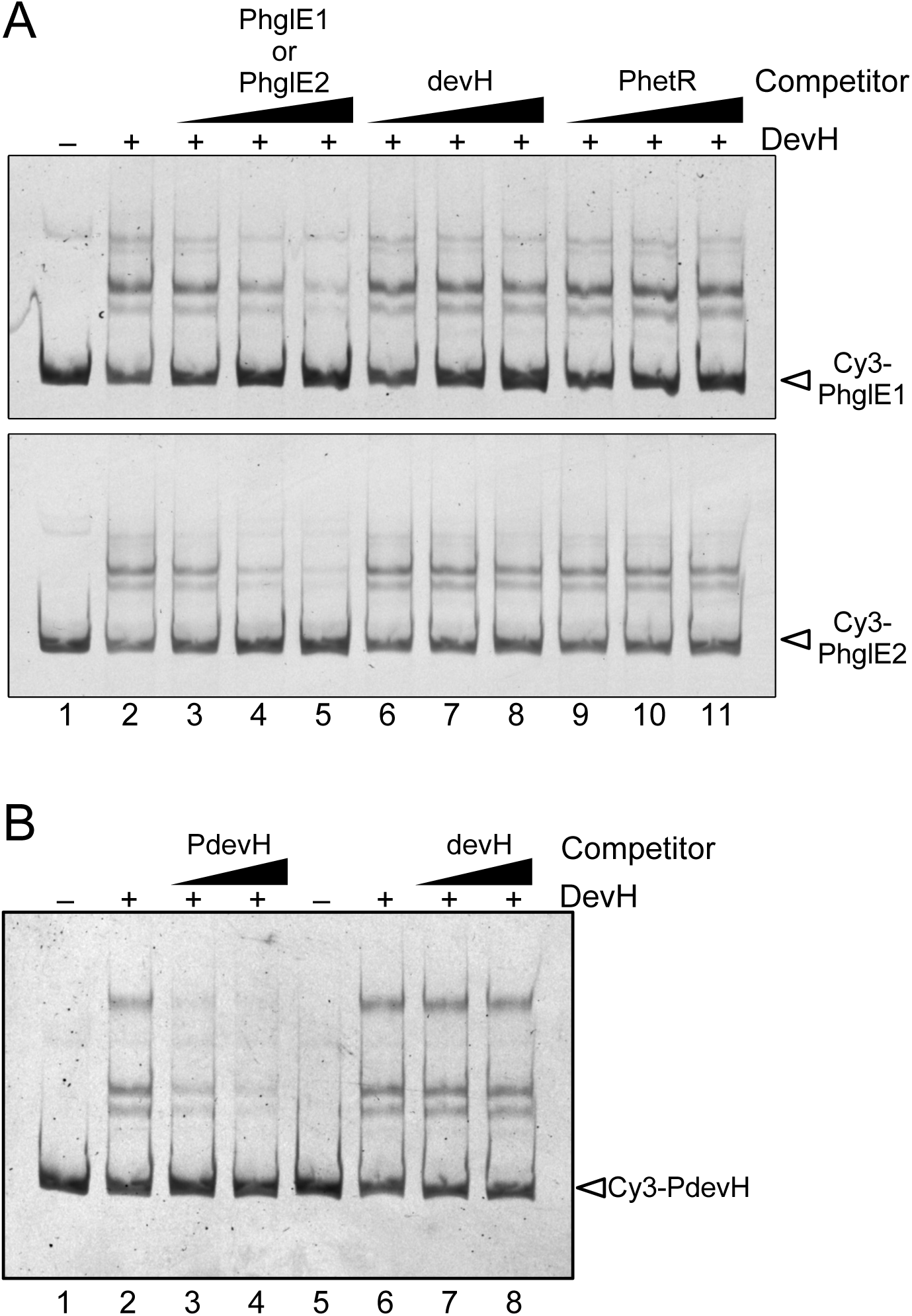
Gel mobility shift assay with a recombinant DevH protein. (A) Interaction between DevH and the proximal *hglE* promoter region (PhglE1) and the distal *hglE* promoter region (PhglE2). DevH (0.1 µM) was mixed with 3 nM Cy3-labeled PhglE1 (above) or PhgelE2 (below). Unlabeled PhglE1 (above) or PheglE2 (below) (lanes 3 to 5), devH (lanes 6 to 8), and PhetR (lanes 9 to 11) were added as competitor DNA at concentrations of 2.5 nM (lanes 3, 6, and 9), 20 nM (lanes 4, 7, and 10), and 50 nM (lanes 5, 8, and 11). Lane 1, DevH was not added. (B) Interaction between DevH and the *devH* promoter region (PdevH). Cy3-labeled PdevH (3 nM) and DevH (0.1 µM) were used for the assays. Unlabeled PdevH (lanes 3 and 4) and devH (lanes 7 and 8) were added as competitor DNA at concentrations of 15 nM (lanes 3 and 7) and 20 nM (lanes 4 and 8). Lanes 1 and 5, DevH was not added.

### *Expression of the* nif *cluster in devHKD under microoxic conditions*

In devHKD, the expression of genes required for creating a microoxic environment within heterocysts was not induced during heterocyst differentiation, which could cause defects in the expression of the *nif* cluster and diazotrophic growth of devHKD. Thus, diazotrophic growth of devHKD under microoxic conditions was examined (see Experimental procedures). An *hglE* mutant, which was unable to perform nitrogen fixation under aerobic conditions (Campbell *et al*., 1997; Fan *et al*., 2005), was used as a positive control. This mutant was able to proliferate using dinitrogen as a nitrogen source under microoxic conditions, similar to WT, but devHKD was not (Fig. 6A). Thus, devHKD is unable to conduct nitrogen fixation even under microoxic conditions.

**Fig. 6.**
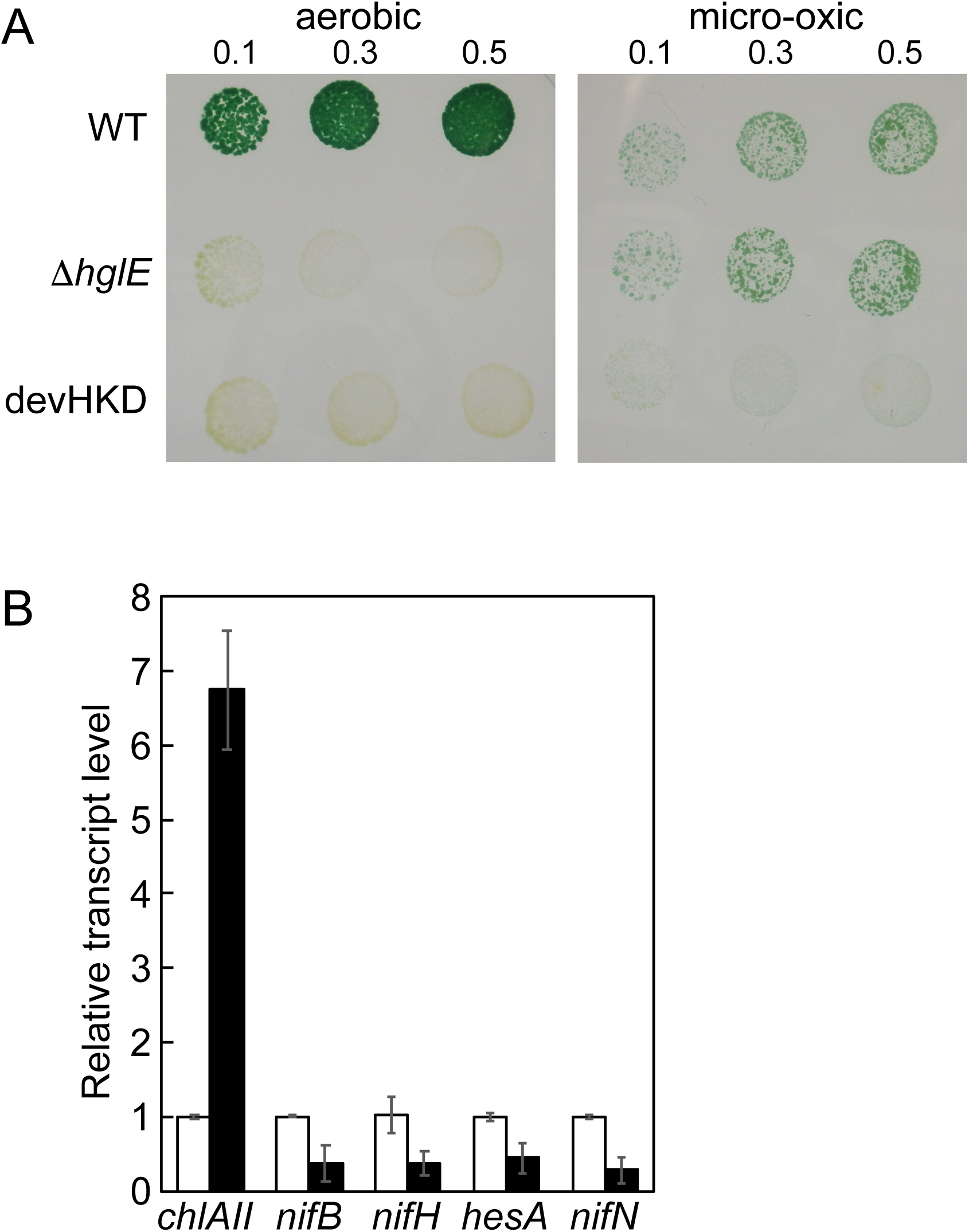
Cultivation of devHKD under microoxic conditions. (A) Cells of WT, an *hglE* mutant strain (Δ*hglE*), and devHKD grown with nitrate were washed with water, the cell densities were adjusted to OD_750_ of 0.1, 0.3, or 0.5, and then aliquots (10 µL) were spotted onto nitrogen-free agar plates. Plates were incubated under aerobic or microoxic conditions for one week. (B) Cells of devHKD depleted of nitrogen for 24 h were spread onto nitrogen-free agar plates and incubated for 18 h under microoxic conditions. The transcript levels under aerobic (white bars) and microoxic (black bars) conditions were determined by qRT-PCR. The qRT-PCR analysis was conducted in duplicate using three independently grown cultures. The transcript level under aerobic conditions was taken as 1.

Expression of the *nif* cluster in devHKD under microoxic conditions was determined. Filaments of devHKD incubated in nitrogen-free liquid medium for 24 h were collected, and then spread on nitrogen-free agar medium followed by incubation under microoxic conditions for 18 h. *chlAII* (all1880) was used as a positive control of microoxic conditions. The *chlAII* expression increases in response to low oxygen under the control of a transcriptional regulator, ChlR, in *Synechocystis* sp. PCC 6803 (Aoki *et al*., 2012) and is upregulated within heterocysts in *Anabaena* PCC 7120 (Ehira, 2013). In this study, the transcript level of *chlAII* increased upon incubation under microoxic conditions, whereas expression of the *nif* cluster under microoxic conditions was lower than that under aerobic ones (Fig. 6B). Thus, the expression of the *nif* cluster is not induced even under microoxic conditions unless DevH is present.

## Discussion

In this study, we demonstrated that DevH regulated gene expression at the later stage of heterocyst differentiation. DevH was shown to be required for inducing the expression of genes encoding oxygen-consuming enzymes and nitrogen fixation genes, which makes it indispensable for the diazotrophic growth of *Anabaena* PCC 7120 (Fig. 7). Because we were unable to inactivate *devH* even under nitrogen-replete conditions, the *devH* expression under nitrogen-deprived conditions was lowered by replacing its promoter with the *gifA* promoter in devHKD (Fig. 1). The *gifA* promoter would be useful for analyzing functions of essential genes under nitrogen-deprived conditions, such as *glnA* and *ntcA*. In addition to regulating heterocyst differentiation, DevH should play a crucial role in vegetative cells. In previous studies by the Curtis’s group, the *devH* mutant strain A57, which expressed DevH proteins with the insertion of 58 amino acids between the C-terminally located helix-turn-helix domain and the C-terminus, was used (Hebbar and Curtis, 2000; Ramírez *et al*., 2005). The DevH protein of A57 would retain some functions that are related to growth under nitrogen-replete conditions.

**Fig. 7.**
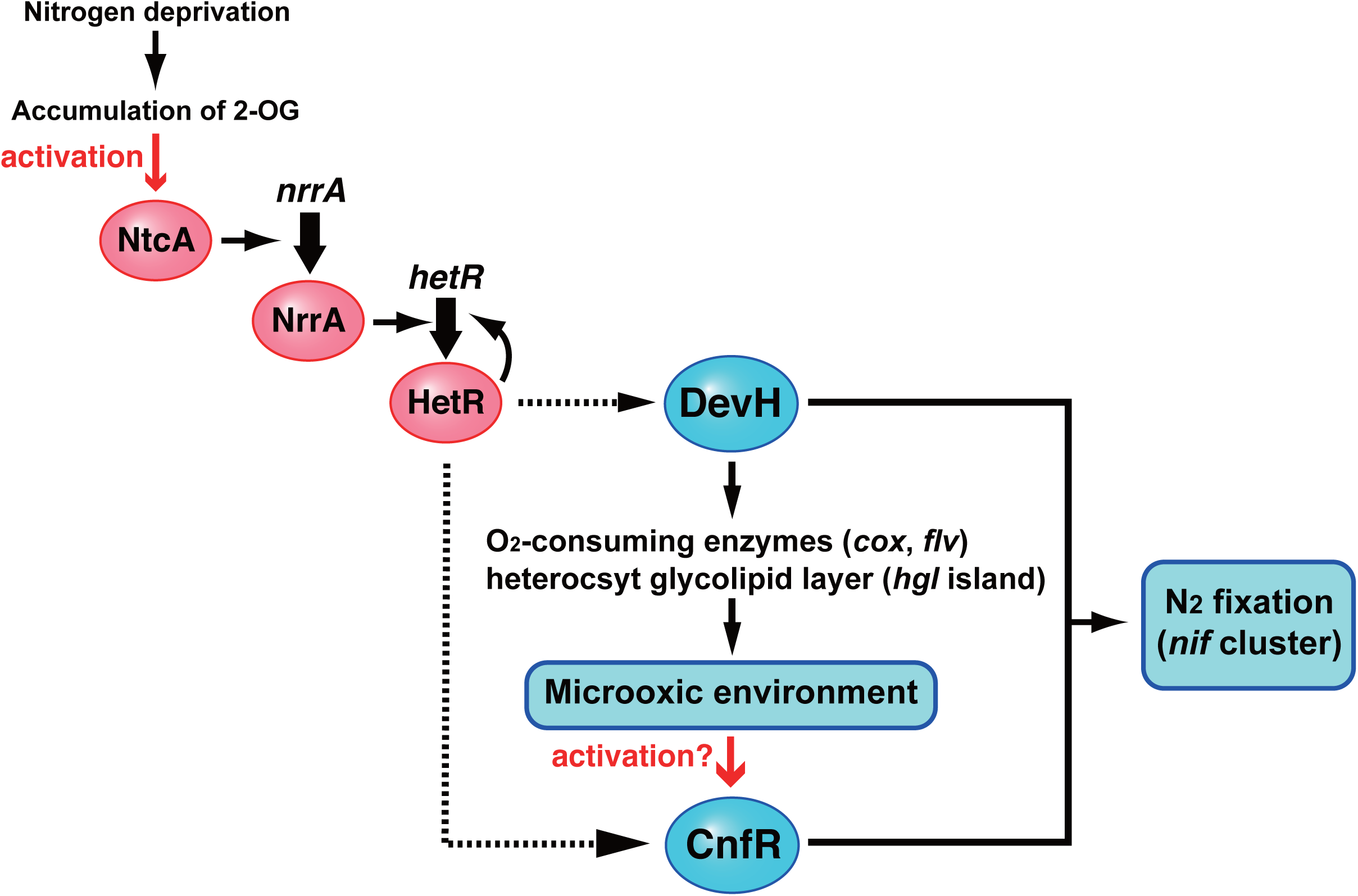
Schematic representation of a regulatory network of heterocyst differentiation in *Anabaena* PCC 7120. NtcA activated by a nitrogen deprivation signal upregulates the *hetR* expression via NrrA. DevH and CnfR, which are under the control of HetR, induces the expression of the *nif* gene cluster. For details, see the text.

Expression of the *nif* cluster was completely abolished in devHKD (Figs. 2 and 3). In the *cnfR* mutant, no induction of *nifB* was observed, but *nifH* was expressed at a lower level than in WT, and *nifN* and *hesA* were expressed at the same level as in WT (Fig. 3), as is the case in *A. variabilis* (Pratte and Thiel, 2016). Expression of the *nif* cluster that is expressed in vegetative cells under anaerobic conditions is solely regulated by CnfR (Tsujimoto *et al*., 2014; Pratte and Thiel, 2016), but the *nif* cluster that is expressed in heterocysts is regulated by CnfR and DevH. Expression of both *cnfR* and *devH* depends on *hetR* in *Anabaena* PCC 7120 (unpublished data) (Hebbar and Curtis, 2000). Dual regulation by CnfR and DevH would ensure the heterocyst-specific expression of nitrogenase and prevent the ectopic expression of nitrogenase in vegetative cells under transient anaerobic conditions (Fig. 7). Moreover, it is noteworthy that the *nif* cluster was not induced even under microoxic conditions in devHKD (Fig. 6). CnfR has a ferredoxin-like domain in its N-terminal region and is supposed to be activated under microoxic conditions (Tsujimoto *et al*., 2016). Because the CnfR level in devHKD was about half of that in WT (Fig. 4), CnfR could induce the *nifB* expression under microoxic conditions even in devHKD if it were activated. When CnfR is constitutively expressed in *Synechocystis* PCC 6803, activation of transcription from the *nifB* promoter by CnfR requires not only microoxic conditions, but also nitrogen-depleted conditions (Tsujimoto *et al*., 2016). Hence, the CnfR activity would be collectively regulated by oxygen levels and another signal such as electron transfer activity, which would be affected by DevH through the regulation of respiration.

Many genes, such as the *hgl* island, the *cox2* and *cox3* operons, *flv3B*, and *cnfR*, identified as the DevH regulon in this study have been shown to also be regulated by NtcA (Flores *et al*., 2019). NtcA is highly expressed in heterocysts (Olmedo-Verd *et al*., 2006), although it is also expressed in vegetative cells and regulates genes involved in functions of vegetative cells (Picossi *et al*., 2014). The NtcA-dependent heterocyst-specific induction is thought to be accomplished by the high level of NtcA and the presence of coactivators, such as PipX, in heterocysts. DevH is likely to be a coactivator of NtcA in the regulation of genes expressed at the later stage of differentiation. Promoters of genes highly expressed in heterocysts are often very complex (Muro-Pastor *et al*., 2009; Camargo *et al*., 2012). Detailed analyses of the promoter regions, for example, NtcA and DevH dependence of promoters and binding sites of NtcA and DevH, would shed light on the mechanisms of dual regulation.

Although the *devH* expression depends on *hetR* (Hebbar and Curtis, 2000), HetR did not bind to the *devH* promoter region (data not shown). We observed that DevH interacted with its own promoter, indicating its autoregulation (Fig. 5). DevH is a member of the CRP family oftranscriptional regulators, in which the cyclic nucleotide-binding domain is located at the N-terminal region, but the cyclic nucleotide-binding domain is absent (Körner *et al*., 2003). The CRP family of transcriptional regulators responds to a wide spectrum of intra- and extracellular signals such as cAMP, anoxia, the redox state, and 2-OG. DevH would be activated by a *hetR*-dependent signal and upregulate its own expression. Identification of a signal that regulates the DevH activity is crucial for understanding of the regulatory mechanism at the later stage of heterocyst differentiation.

## Experimental procedures

### Bacterial strains and culture conditions

*Anabaena* PCC 7120 and its derivatives were grown in ammonium-containing BG-11 medium buffered with 20 mM 2-[4-(2-hydroxyethyl)-1-piperazinyl] ethanesulfonic acid-NaOH (pH 7.5) (Rippka *et al*., 1979), in which 17.6 mM NaNO_3_ included in the original BG-11 medium was replaced with 5 mM NH_4_Cl. The conditions of growth and heterocyst induction were as previously described (Ehira and Ohmori, 2006a). For growth under microoxic conditions, agar plates of combined nitrogen-free BG-11 medium were incubated in a sealed hybridization bag (Hybri-Bag Soft; Cosmo Bio, Tokyo Japan) with an oxygen absorber-CO_2_ generator (AnaeroPack-Anaero; Mitsubishi Gas Chemical Company, Tokyo, Japan) under continuous illumination at 10 µmol photons m^−2^ s^−1^.

### Mutant construction

The primers listed in Table 1 were designed based on genome data from CyanoBase (Fujisawa *et al*., 2017). Upstream and coding regions of the *devH* gene and the promoter region of the *gifA* gene were amplified by PCR with the primer pairs devH-5F3 and devH-5R, devH-3F3 and devH-3R2, and PgifA-F2 and PgifA-R2, respectively. The *devH* upstream region was cloned between the SacI and BamHI sites of pBluescript II KS^+^ (Agilent Technologies), and then the *devH* coding region was cloned between the SalI and XhoI sites. The *gifA* promoter region was inserted between the upstream and coding regions of *devH* using the BamHI and SalI sites, and a spectinomycin/streptomycin resistance cassette derived from pDW9 (Golden and Wiest, 1988) was inserted between the *devH* upstream region and the *gifA* promoter region. The resultant DNA construct was excised as SacI-XhoI fragments and cloned into the same sites of pRL271 (Black *et al*., 1993) to create pPgifA-devHS. Upstream and downstream regions of the *cnfR* gene were amplified by PCR with the primer pairs 2512-5F and 2512-5R, and 2512-3F and 2512-3R, respectively. The upstream region was cloned between the SacI and BamHI sites of pBluescript II KS^+^, and then the downstream region was cloned between the BamHI and XhoI sites. A kanamycin/neomycin resistance cassette derived from pRL161 (Elhai and Wolk, 1988) was inserted between the upstream and downstream regions. The resultant DNA construct was excised as SacI-XhoI fragments and cloned into the same sites of pRL271 to create pR2512K. A plasmid pRhglES was created as pR2512K using the primer pairs hglE-5F and hglE-5R, and hglE-3F and hglE-3R, and the spectinomycin/streptomycin resistance cassette. pPgifA-devHS, pR2512K, and pRhglES were transferred by conjugation into *Anabaena* PCC 7120 in accordance with the method of Elhai et al. (1997), and double recombinants were selected for resistance to spectinomycin/streptomycin or neomycin and sucrose. The completion of segregation was confirmed by PCR, and the completely segregated clones devHKD, DRcnfRK, and DRhglES were used in this study.

### RNA extraction and qRT-PCR analysis

Total RNA extraction and qRT-PCR analysis were conducted as described previously (Ehira and Miyazaki, 2015). The primers used in the qRT-PCR are listed in Table 1. Transcript levels were normalized against the level of 16S rRNA and determined using three biologically independent samples.

### RNA sequencing analysis

RNAs for RNA sequencing analysis were extracted as described above and further purified by ultracentrifugation. Library construction for RNA sequencing analysis was conducted as described previously (Shimmori *et al*., 2018). Two biological replicates were made for each condition. The average number of raw read pairs per sample was 9.12 million. The reads were trimmed using CLC Genomics Workbench ver. 11.0 with the following parameters; Phred quality score >25; ambiguous nucleotides allowed: 1; automatic read-through adaptor trimming: yes; removing the terminal 15 nucleotides from the 5’ end and 2 nucleotides from the 3’ end; and removing truncated reads of less than 30 nucleotides in length. Trimmed reads were mapped to the all genes in *Anabaena* PCC 7120 (heterotypic synonym, *Nostoc* sp. PCC 7120: accession number: BA000019.2) using CLC Genomics Workbench ver. 11.0 (Qiagen) with the following parameters; match score: 1; mismatch cost: 2; indel cost: 3; length fraction: 0.7; similarity fraction: 0.9; and maximum number of hits for a read: 1. For the identification of differentially expressed genes between the wild-type and the devHKD cells, the edgeR package version 3.4.0 with the exact test and trimmed mean of M-values (TMM) normalization method implemented in the ‘Empirical analysis of DGE’ algorithm in CLC Genomics Workbench was used with following parameters; Total count filter cut off : 10.0; Estimate tagwise dispersions: yes; FDR corrected: yes. The Detailed parameters values used in the algorithm was enclosed in the manufacturer’s product information. Genes with |log2 fold change| > 1 and FDR <0.05 were considered to be differentially expressed between wild-type and mutant cells. Original sequence reads were deposited in the DRA/SRA database with the following accession numbers (DRR215823– DRR215826). The accession number of BioProject was PRJDB9455.

### Expression and purification of DevH recombinant proteins

DevH protein of *Anabaena* PCC 7120 was expressed as a fusion protein with a solubilizing tag, ProS2. The *devH* coding region amplified by PCR using the primer pair NHisdevH-F1 and NHisdevH-R1 was cloned between the NdeI and SalI sites of the expression vector pColdProS2 (Takara Bio, Shiga, Japan) to construct pCProS2DevH. Expression and purification of DevH recombinant proteins were conducted as described previously (Ehira and Ohmori, 2012).

### Gel mobility shift assay

Gel mobility shift assay was performed using recombinant DevH proteins and 3 nM Cy3-labeled DNA probes in 20 µl of binding buffer [20 mM Tris-HCl (pH 7.0), 10 mM MgCl_2_, 40 mg L^−1^ bovine serum albumin, 1 mM dithiothreitol, 5% glycerol]. The mixtures were incubated for 30 min at room temperature and then subjected to electrophoresis on a native 5% polyacrylamide gel. DNA probes were visualized with the FLA9000 image scanner (FUJIFILM, Tokyo, Japan). Cy3-labeled DNA probes were prepared as follows; Each promoter region was amplified by PCR using the primers listed in Table 1, and the PCR products were cloned into the EcoRV site of pBluescript II KS^+^. The resultant plasmids were used for PCR as templates with a Cy3-labeled M13-F primer.

## Acknowledgments

This work was supported by the Japan Society for the Promotion of Science (JSPS) [Challenging Research Exploratory 19K22290], the Iwatani Naoji Foundation, the Nagase Science and Technology Foundation, and Cooperative Research Grant of the Genome Research for BioResource from NODAI Genome Research Center, Tokyo University of Agriculture to SE.

## Author contributions

SE conceived and designed the study; YKu, YKo, YKa and SE acquired and analyzed the data; SE and YKa wrote the manuscript, and all authors edited the manuscript.

